# Mucosal administration of anti-bacterial antibody provides long-term cross-protection against *Pseudomonas aeruginosa* respiratory infection

**DOI:** 10.1101/2022.07.04.498699

**Authors:** Aubin Pitiot, Marion Ferreira, Christelle Parent, Chloé Boisseau, Mélanie Cortes, Laura Bouvart, Christophe Paget, Nathalie Heuzé-Vourc’h, Thomas Sécher

## Abstract

Bacterial respiratory infections, associated with acute, sometimes recurrent infections and with chronic respiratory diseases, are a major threat for human health. Mucosal administration of therapeutic antibodies (Ab), using the airways as a delivery route, has a tremendous opportunity to benefit to patients with respiratory infections, with remarkable preclinical achievements in both viral and bacterial respiratory infection models and ongoing clinical developments. The primary mode of action of anti-infective Ab delivered through the airways is pathogen neutralization and to a lesser extent, Fc-mediated direct recruitment of immune effectors to facilitate their elimination. Using a mouse model of acute pneumonia induced by *P. aeruginosa*, a bacterium frequently associated with multidrug resistance and a high rate of recurrence, we characterized an immunomodulatory mode of action of anti-bacterial Ab. Beyond the rapid and efficient containment of the primary infection, the anti-infective Ab delivered through the airways harnessed adaptive immunity to provide a long-term response, preventing from a secondary pathogen infection. This effect is specific and dependent on the Ab dose, intensity of infection and antigen expression by the pathogen upon primary infection. As shown by adoptive transfer experiments, it is mediated by a sustained and protective humoral immune response. Interestingly, the long-lasting response protected partially against secondary infections due to heterologous *P. aeruginosa* strains. Overall, our findings suggest that mucosal delivery of Ab through the airways offers a dual advantage: a rapid onset of action to neutralize respiratory bacteria and a long-term protection against secondary infections, thereby opening novel perspectives for the development of anti-infective antibody delivered to the lung mucosa, to treat respiratory infections.

## Introduction

Since the advent of monoclonal antibody technology, the use of antibodies (Ab) has proven to be successful for the treatment or prevention of a variety of disorders. Ab have radically changed the management and treatment of fatal diseases. The success of Ab is based on their high specificity, predictable toxicity and unique pharmacological profiles, giving them advantages over small molecule drugs. Beyond cancer and inflammatory diseases, Ab are, now, considered as potential game changers for the management of infectious diseases. Actually, four anti-infective Ab are already approved, targeting either bacterial or viral pathogens and several other entities already reached clinical trials and may enrich the therapeutic armamentarium against infectious diseases ^1, 2^. Overall, anti-infective Ab therapy offers prophylactic or therapeutic advances against infections when vaccines and/or conventional drugs are neither available nor efficacious, by overriding an immune system that is unresponsive to immunization or countering pathogen resistance.

Anti-infective Ab have multifaceted mechanisms of action. Few of them are designed to inhibit pathogen toxin, while the majority of Ab recognize cell surface antigen, through their antigen-binding (Fab) domain, thereby preventing the attachment, entry or the production of virulence factors into the host cell. Although anti-infective Ab are primarily neutralizing, killing of the pathogen may also be facilitated by effector functions, including antibody-dependent cellular phagocytosis/cytotoxicity (ADCP/ADCC), complement-dependent cytotoxicity (CDC), that depend on the interaction between the crystallizable fragment (Fc) domain and Fcγ receptors (FcγR) or enzyme cascade of the complement.

Accumulating evidences highlight a novel function of Ab, which can engage and promote endogenous innate and adaptive immune systems to induce a long-lasting protection ^3^. This has been widely exemplified at the preclinical and clinical level, with antitumor- and anti-viral Ab. For instance, patients treated with anti-MUC1 or anti-HER2 Ab generate anti-MUC-1 and HER2-specific T cell responses ^4, 5^. Similarly, a single course of treatment with anti-CD20 Ab results in long-lasting tumor surveillance in patients with lymphoma ^6^, in whom an anti-idiotype T cell response against lymphoma antigens was detected ^7, 8^. Ab-mediated immune-stimulating response has also been illustrated during viral infections ^9, 10^. A notable common characteristic, in both disease contexts, is the importance of the Fc domain, which connects specific recognition of tumor or viral antigens with immune cells mediating antibody-dependent cellular cytotoxicity or phagocytosis (ADCC or ADCP) through the engagement of FcγR. Immune complexes (IC), formed between therapeutic Ab and tumor or viral particles infected, will promote antigen-presenting cells stimulation, maturation and presentation of pathogenic antigens to T cells, eventually leading to long-term immune memory response ^3, 11^. The inhibition of regulatory T cell expansion was mandatory for the induction of these protective responses ^12, 13^. Interestingly, while Ab-mediated long-term anti-tumor protection was solely associated with cellular responses ^14^, anti-viral Ab prompted both host cellular and humoral immunity ^15, 16^, essential to long-term containment of viremia. Although the mechanisms underlying Ab-mediated long-lasting response are not identical across disease contexts, taken as a whole, those findings bring a new concept in Ab immunotherapy. Here, we speculated that Ab-mediated long-term protection may apply to neutralizing anti-bacterial Ab and benefit to the treatment of respiratory infections, in particular those due to pathogens associated to persistent chronic or recurrent acute respiratory infections, like the ESKAPE bacteria (*Enterococcus faecium, Staphylococcus aureus, Klebsiella pneumoniae, Acinetobacter baumannii, Pseudomonas aeruginosa* and *Enterobacter* spp.), which are often multidrug resistant and a major threat for human health.

In respiratory medicine, drugs can be delivered parenterally or locally, using the airways as a direct portal to the lungs. Although, intravenous (i.v) injection is the conventional route for anti-infective Ab, it is not optimal for the delivery of full-length Ab to the airways, where pathogens mainly infect, propagate and disseminate. Indeed, the transport of Ab across the lung epithelial layer is relatively inefficient in either direction ^17-19^. In contrast, others and we have shown that airway administration of Ab conferred greater protection than parenteral administration ^20-23^. Airway delivery favors a rapid onset of action, resulting in the efficient neutralization of pathogens ^22, 23^ and the pharmacokinetics (PK) profile is favorable: Ab accumulate at higher concentrations in the airways than after i.v infusions and pass slowly and in low amounts from the airways into the circulation ^19, 22^. In addition, the respiratory mucosa, which is directly accessible through the airways, provides a specific immune environment, favorable for establishing long-term protection. Indeed, comparative studies have shown that mucosal routes provide superior efficacy in the induction of mucosal immune responses as compared to systemic administration ^24^. In particular, higher protection was obtained, at local or even remote sites, against airway pathogens when immunization was applied by the mucosal route ^25, 26^. Here, we speculated that mucosal delivery of Ab through the airways is well-suited to achieve both direct control and long-term containment of bacterial respiratory pathogens.

Using an acute model of respiratory infection, due to *Pseudomonas aeruginosa*, a bacterium with a frequent multi-resistance phenotype and high-rate of recurrence, we found that mucosal delivery of neutralizing Ab provided both an optimal defense against primary infection and frontline immunity for the remote protection against subsequent infections. Ab-mediated long-lasting protection was dependent on the size of the inoculum, the dose of Ab and the presence of the cognate antigen during the primary infection. Remarkably, the immune response was sufficient to promote partial heterologous protection against *P. aeruginosa* strains from different serotypes. The long-lasting response of mucosal Ab in bacterial respiratory infection relies, at least partly, on the establishment of a sustained and protective humoral immune response.

## Results

### Prophylactic and therapeutic mucosal administration of anti-bacterial antibody, through the airways, mediates long-term protection against *P. aeruginosa*

As previously demonstrated, mAb166, a murine IgG2b recognizing PcrV – an antigen belonging to the type 3 secretion system (T3SS), protects mice from a lethal pulmonary infection with *P. aeruginosa* ^22^ after airway delivery. Here, we investigated whether the rescued animals were protected from subsequent lung infection.

Mice were treated with 100µg of inhaled mAb166 either prior or after being inoculated intrapulmonary with a lethal dose of *P. aeruginosa* PA103 strain (Figures 1A and 1C) to mimic both a prophylactic and therapeutic Ab intervention. After 30 days, the surviving mice were re-infected with the same lethal dose of PA103 (as for the primary infection), without additional mAb166 treatment, and at a time where mAb166 was no longer detectable in the serum and broncho-alveolar lavage fluid (BALF) (Supplementary Figure 1 and ^22^). It is noteworthy that the prophylactic intervention afforded 80% to 90% protection (Figure 1B) after the primary infection, while the therapeutic one rescued 60% to 80% animals (Figure 1D). After the secondary infection, all the animals that previously survived from the primary infection resisted the lethal-dose infection (Figures 1 B and D). This beneficial effect of the immunotherapy was associated with an improved control of the bacterial load in both lung and BAL after the primary and the secondary infection (Figure 1E)

**Figure 1:**
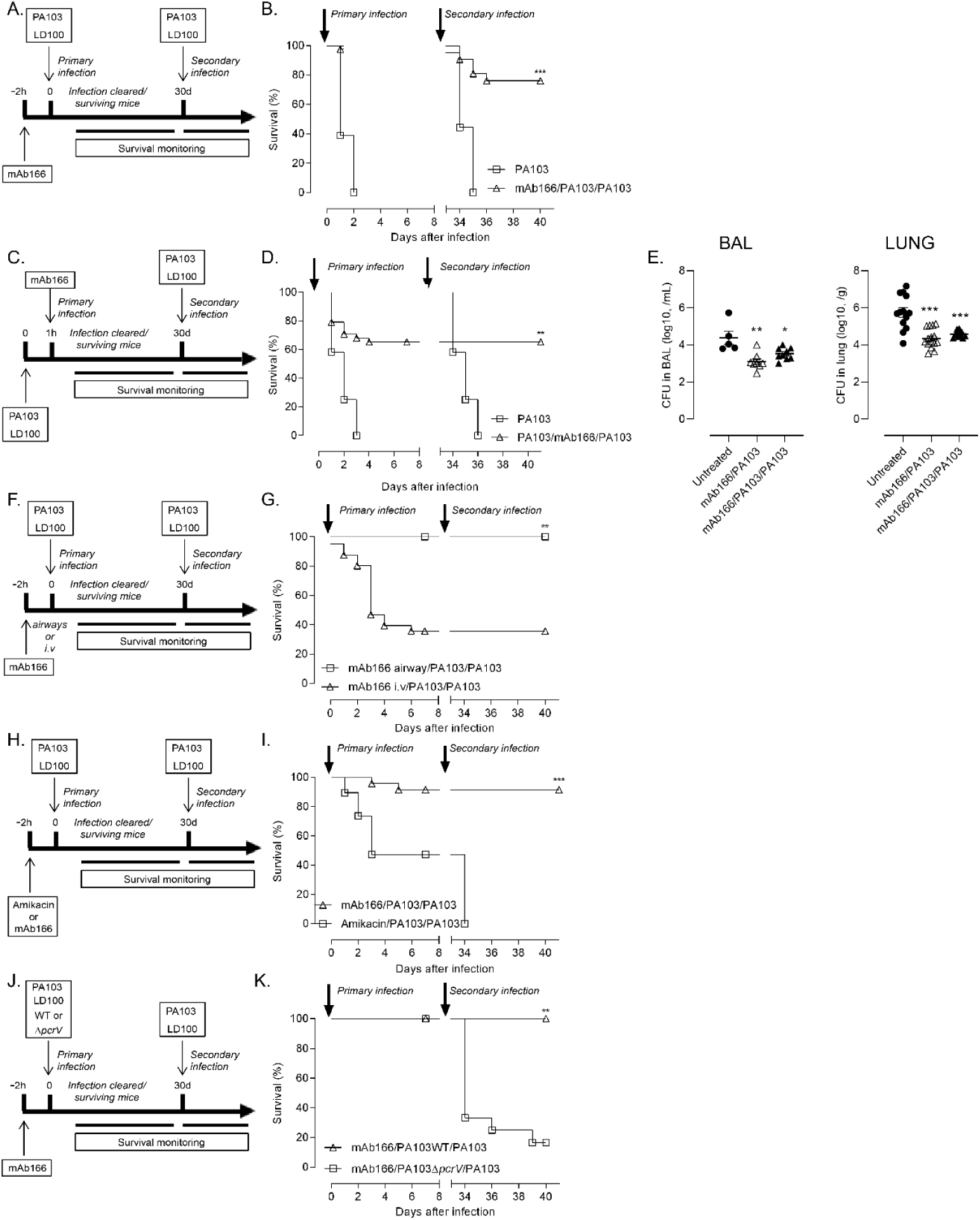
Prophylactic and therapeutic mucosal administration of anti-bacterial antibody, through the airways, mediates long-term protection against P. aeruginosa. (A) C57/BL6jRj (B6) mice were treated or not with 100 µg of mAb166 via the pulmonary (airway) route 2 hours before being infected by the orotracheal instillation of 40 µL of *P. aeruginosa* strain PA103 (5×10^5^ cfu / primary infection). Surviving animals were challenged, 30 days later (secondary infection) by the orotracheal instillation of 40 µL of *P. aeruginosa* strain PA103 (5×10^5^ cfu) without additional treatment. (B) Survival was monitored over the entire period. The results correspond to 6 pooled, independent experiments (n=27-43 mice per group), ***: *p<*0.001 with a log-rank test. (C) B6 mice were infected by the orotracheal instillation of 40 µL of *P. aeruginosa* strain PA103 (3×10^5^ cfu / primary infection) and 1 hour later were treated or not with 100 µg of mAb166 via the pulmonary (airway) route. Surviving animals were challenged, 30 days later (secondary infection) by the orotracheal instillation of 40 µL of *P. aeruginosa* strain PA103 (3×10^5^ cfu) without additional treatment. (D) Survival was monitored over the entire period. The results correspond to 6 pooled, independent experiments (n=36-72 mice per group), **: *p<*0.01 with a log-rank test. (E) The bacterial load (cfu) in the broncho-alveolar lavage (BAL) and the lungs was measured 24 hours after the primary infection or after the secondary infection. The data are quoted as the mean values ± SEM. The results correspond to 4 pooled, independent experiments (n=5-15 mice per group), *: *p<*0.05; **: *p<*0.01; ***: *p*<0.001 with a *t* test test comparing with untreated groups. (F) B6 mice were treated with 100 µg of mAb166 via the pulmonary (airway) or intravenous (i.v.) 2 hours before being infected as described in (A). (G) Survival was monitored over the entire period. The results correspond to 2 pooled, independent experiments (n=8-27 mice per group), **: *p<*0.01 with a log-rank test. (H) B6 mice were treated with 100µg of mAb166 or 25mg/kg of Amikacin and were infected as described in (A). (I) Survival was monitored over the entire period. The results correspond to 2 pooled, independent experiments (n=19-24 mice per group), ***: *p<*0.001 with a log-rank test. (J) B6 mice were treated with 100µg of mAb166 and infected with PA103 WT or with PA103Δ*pcrV* as described in (A). (K) Survival was monitored over the entire period. The results correspond to 2 pooled, independent experiments (n=15-24 mice per group) **: *p<*0.01 with a log-rank test.

In comparison to systemic administration (i.v) (Figure 1F), which is the conventional delivery route for anti-infective Ab, the overall response to repeated bacterial infections was significantly better if the mAb166 was delivered through the airways upon the primary infection (Figure 1G). It is noteworthy that the benefit of mucosal delivery is mainly attributable to a better control of primary infection, since the animals exhibited similar survival rates after the secondary infection, whatever the Ab route of administration. Overall, this highlights the superiority of the airways for the delivery of anti-infective Ab to contain pathogens after primary and subsequent infections.

To investigate the specificity of this long-lasting response, we first assessed the capacity of standard anti-*P. aeruginosa* antibiotic therapy (amikacin) to protect animals from multiple infections. None of the animals that were rescued from the primary infection after airway administration of amikacin (Figure 1H), survived a secondary infection, without additional treatment (Figure 1I). Secondly, we infected mice with an isogenic PA103 mutant deprived of *pcrV* expression, PA103Δ*pcrV* (Figure 1J). This attenuated *P. aeruginosa* strain, caused morbidity (Supplementary figure 2B) but not mortality in animals (Supplementary figure 2C). Interestingly, the animals primarily infected with PA103Δ*pcrV* and treated with mAb166, were not able to survive a second infection with the wild-type PA103 (Figure 1K). Overall, these results indicate that the induction of a long-term protection against *P. aeruginosa* by Ab delivered through the airways depends on the antibody binding to its target antigen, as previously shown with anti-tumor antibody ^11^.

### Antibody-mediated long-term protection against *P. aeruginosa* depends on a subtle balance between bacteria inoculum and Ab dose

To further characterize long-term protection induced by prior mucosal Ab treatment, we treated animals with a suboptimal quantity of mAb166 (50µg) (Figure 2A), leading to 40-50% of protection after the primary infection (Figure 2B). Interestingly, this was not sufficient to protect all the rescued animals from a second infection (Figure 2B). Similarly, mice infected with a non-lethal dose of PA103 (5.10^4^cfu) (Figure 2C) which caused morbidity (Supplementary Figure 3C) but not mortality (Supplementary Figure 3C), were not protected from a second infection with a lethal dose of PA103 (Figure 2B). We did not observe any protection against PA103 secondary infection when mice were previously treated by mucosal prophylactic delivery of mAb166-PA103 immune complexes (data not shown). Taken together, these results demonstrated that features of active infection and optimal mucosal treatment are essential to the generation of a long-term immunity against respiratory pathogens.

**Figure 2:**
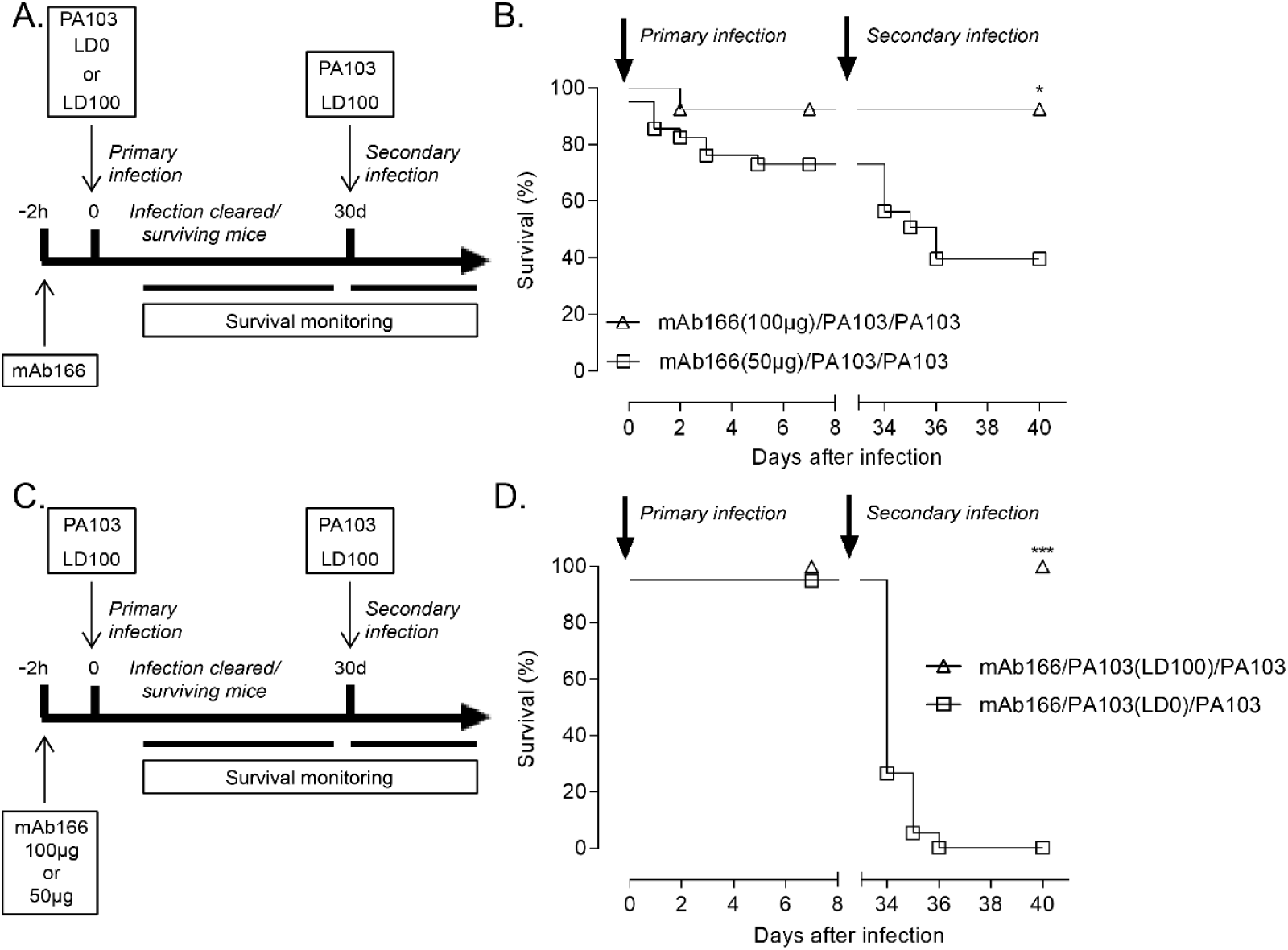
Antibody-mediated long-term protection against P. aeruginosa depends on a subtle balance between bacteria inoculum and Ab dose. (A) B6 mice were treated with 100µg or 50µg of mAb166 and infected as described in Figure 1A. (B) Survival was monitored over the entire period. The results correspond to 4 pooled, independent experiments (n=27-32 mice per group), *: *p<*0.05 with a log-rank test. (C) B6 mice were treated with 100µg of mAb166 and infected with a lethal dose (5×10^5^ cfu / primary infection) or a non-lethal dose (5×10^4^ cfu / primary infection) as described in Figure 1A. (D) Survival was monitored over the entire period. The results correspond to 5 pooled, independent experiments (n=16-30 mice per group), ***: *p<*0.001 with a log-rank test.

### Long-term protection is associated with improved lung inflammatory response against *P. aeruginosa*

To gain first insight into the mechanisms accounting for long-term lung protection during *P. aeruginosa* infection, we quantified, in the airways and lung tissue, the recruitment of neutrophils – the prototypal leukocytes associated with the control of *P. aeruginosa* infections ^27^. After 4h, we observed that they were significantly more neutrophils in both BALF and lungs of the protected animals as compared to the susceptible controls (Figures 3A and B). This was accompanied by a significant reduction of PA103 load (Figures 3C and D). At 24h after secondary infection, a time at which they were less bacteria in the lung of mice, there were no longer differences in the number of airway’s neutrophils (Figures 3A and B). These data suggest that long-term lung protection generated by mucosally-administered Ab remodel lung immune response to improve neutrophil recruitment, which in turn achieves a better control of *P. aeruginosa*.

**Figure 3:**
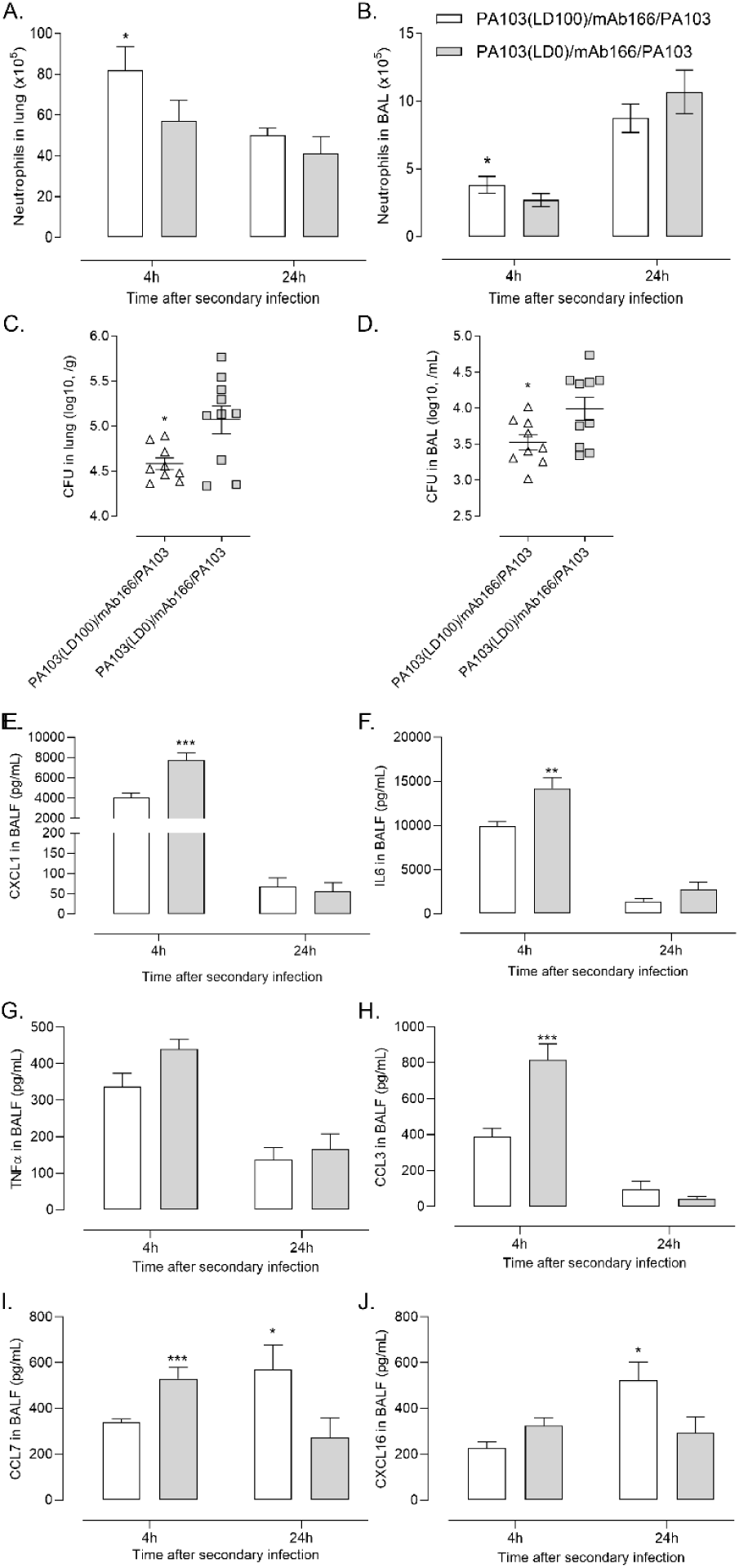
Long-term protection is associated with improved lung inflammatory response against *P. aeruginosa*. B6 mice were treated/infected as described in Figure 1C. (B) Neutrophils (CD45+ CD11b+ SiglecF-Ly6G+ cells) number in lung (A) and BAL (B) were determined 4 and 24 hours after the secondary infection. The data are quoted as the mean values ± SEM. The results correspond to 3 pooled, independent experiments (n=13 mice per group), *: *p<*0.05 with *t* test. Bacterial load in lung (C) and BAL (B) were determined 4 hours after the secondary infection. The data are quoted as the mean values ± SEM. The results correspond to 2 pooled, independent experiments (n=10 mice per group), *: *p<*0.05 with *t* test. At 4 and 24 hours after the secondary infection, BALF were analyzed by Multiplex assay for the production of (E) CXCL1, (F) IL-6, (G) TNFα, (H) CCL3, (I) CCL7, (J) CXCL16. The data are quoted as the mean values ± SEM. The results correspond to 3 pooled, independent experiments (n=13 mice per group), *: *: *p<*0.05; **: *p<*0.01; ***: *p*<0.001 with a *t* test.

While neutrophilic-dependent immune response is necessary for bacterial clearance, a protracting and non-resolving inflammatory response is associated with poor host outcomes in *P. aeruginosa* acute lung infection model ^28^. Therefore, we investigated pro-inflammatory cytokines and chemokines in BALF upon *P. aeruginosa* secondary infection. Interestingly, the expression of the innate immunity mediators CXCL1, IL6, CCL3 and to a lesser extent TNF (Figures 3E-H) was significantly reduced in the BALF of mice with protective immunity within 4 hours after the challenge. Despite an increased neutrophils influx, dampening of proinflammatory pathways may mediate the early resolution of the acute inflammatory response. At later time-point, the levels of CXCL16 and CCL7, two chemokines associated with the recruitment and activation of cells associated with the adaptive immunity, were significantly increased in mice protected from secondary infection (Figures 3I and 3J). Overall, our data suggested that protection to *P. aeruginosa* secondary may be attributable toa better defense of the mucosal surface, which is promoted by an improved recruitment and activity of neutrophils as well as partners from the adaptive immunity.

### Mucosal administration of mAb166 leads to the development of a sustained and protective humoral immune response against *P. aeruginosa*

The enhanced lung bacterial clearance after the secondary infection might be due to improved uptake/bactericidal features of recruited neutrophils. This may be due by a better bacterial opsonization afforded by circulating anti-*P. aeruginosa* IgG. Indeed, humoral immunity is the main specific immune response to protect against extracellular pathogenic bacteria ^29, 30^.

To delineate the immune mechanisms of long-term protection against bacterial pneumonia, we analyzed the primary as well as recall humoral response against *P. aeruginosa*. We observed a significant enrichment of anti-PA103 IgG in the serum of mice infected with a lethal dose of PA103 and treated with an optimal quantity of mAb166 as compared to the ones infected with a non-lethal dose (Figure 4A), the Δ*pcrV* mutant or animals treated with a suboptimal quantity of mAb166 (Supplementary Figure 4A). Moreover, under protective conditions, we observed a rapid and robust induction of the humoral response after the secondary infection (Figures 4B). Interestingly, comparing before and after the secondary infection, we showed a significant enrichment of anti-PA103 IgG in the serum only in protected mice as compared to susceptible controls suggesting the induction of a memory humoral response (Figure 4C). We next addressed the frequencies of CD19+ B cells in the spleen, as a surrogate of systemic and local humoral responses, at one day after the secondary infection. The number of systemic B cells was enhanced in mice which developed a long-term protection (Figures 4D and 4E) with a significant proportion of them with a MZ phenotype (CD21^h^ IgM^h^, CD19^+^) (data not shown), which are critical to the antibody responses ^31^. It is noteworthy that similar results were obtained in the prophylactic intervention model (Supplementary Figure 4). We next investigated whether *P. aeruginosa* prior exposure established such humoral response within the airways and observed while IgG titers in BALF were present in a lesser extent as compared in serum (Figure 4F), they were significantly enriched in protected mice (Figure 4G). This was associated with an enrichment of CD19+ B cells in the lung parenchyma (Figures 4H and I) as well as an increased expression of BAFF – which is known as a crucial cytokine for B cell activation and maturation ^32^ – 24h after the secondary infection (Figure 4J).

**Figure 4:**
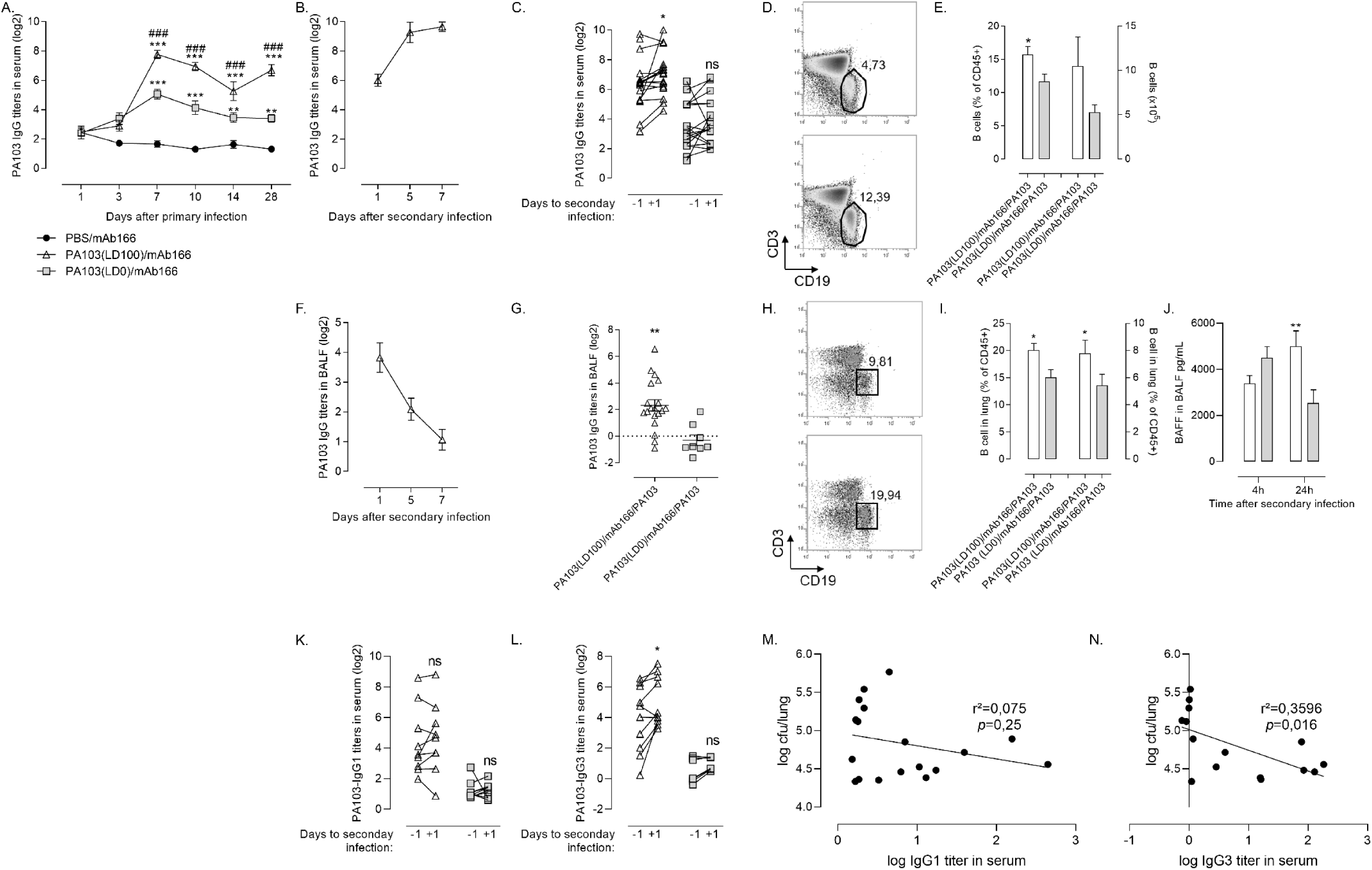
Mucosal administration of mAb166 leads to the development of a sustained and protective humoral immune response against *P. aeruginosa*. B6 mice were treated/infected as described in Figure 1C. (A) The concentration of total anti-PA103 IgG in serum was determined by ELISA at days 1, 3, 7, 10, 14, 28 after the primary infection. The data are quoted as the mean values ± SEM. The results correspond to 7 pooled, independent experiments (n=10-33 mice per time-points), ###: *p<*0.001 with two-way ANOVA followed by a Bonferonni’s post-test, comparing PA103(3.10^5^)/mAb166 with PA103(5.10^4^)/mAb166. **: *p<*0.01; ***: *p<*0.001 with two-way ANOVA followed by a Bonferonni’s post-test, comparing with PBS controls. (B) The concentration of total anti-PA103 IgG in serum was determined by ELISA at days 1, 5 and 7 after the secondary infection. The data are quoted as the mean values ± SEM. The results correspond to 3 pooled, independent experiments (n=10 mice per time-points). (C) The concentration of total anti-PA103 IgG in serum was determined by ELISA at days -1, +1 after the secondary infection. The data are quoted as individual values. The results correspond to 4 pooled, independent experiments (n=17-19 mice per group), *: *p<*0.05; with a paired t-test. 24 hours after the secondary infection, (D) representative dot plot depicting CD19+ B cells in the spleen, (E) frequency and total number of splenic CD19+ B cells (CD45+ CD3-CD19+ cells). The data are quoted as the paired mean values ± SEM. The results correspond to 2 pooled, independent experiments (n=10 mice per time-points), *: *p<*0.05; with a *t* test. (F) The concentration of total anti-PA103 IgG in BALF was determined by ELISA at days 1, 5 and 7 after the secondary infection. The data are quoted as the mean values ± SEM. The results correspond to 6 pooled, independent experiments (n=8-18 mice per time-points). (G) the concentration of total anti-PA103 IgG in BALF was determined by ELISA at day 1 after the secondary infection. The data are quoted as the mean values ± SEM. The results correspond to 3 pooled, independent experiments (n=8-19 mice per time-points). **: *p<*0.01; with a *t* test. 24 hours after the secondary infection, (H) representative dot plot depicting CD19+ B cells in the lung, (I) total number of lung CD19+ B cells and (J) BAFF expression in the BALF. The data are quoted as the paired mean values ± SEM. The results correspond to 2 pooled, independent experiments (n=10 mice per time-points), *: *p<*0.05; **: p<0.01 with a *t* test. The concentration of anti-PA103 IgG1 (K) and IgG3 (L) in serum was determined by ELISA at days -1, +1 after the secondary infection. Correlation analysis of PA103 burden (depicted in Figure 3C) in lung and IgG1 (M) or IgG3 (N) in the serum. The data are quoted as individual values. The results correspond to 4 pooled, independent experiments (n=17-19 mice per group), *: *p<*0.05; with a paired t-test.

We next assessed the relative amount of different anti-PA103 IgG subclasses present in animals exhibiting different ability to limit the extent of the secondary infection. IgG1 and IgG3 were the most abundant isotypes found in the serum and BALF (data not shown) after *P. aeruginosa* infection. We observed a significant enrichment of IgG3 on the contrary to IgG1 in the serum of the mice resistant to the secondary infection, as compared to the susceptible animals (Figures 4K and 4L). Indeed, a formal correlation analysis of bacterial burden in the lung with IgG titers in the serum showed that the presence of IgG3 correlated with a better bacterial control, as compared to IgG1 after PA103 secondary infection (Figures 4M and 4N). Overall, our data support the idea that the mice infected and treated with mAb166, under optimal conditions, resist re-infection by PA103 due to an efficient anti-bacterial humoral immune response, which is recalled after the second bacterial infection.

To investigate whether this protracted humoral anti-PA103 response was protective on its own, we collected sera from animals that were either protected from a secondary infection or not and uninfected/mAb treated controls mice. Sera were decomplemented and administered intraperitoneally to naive mice (Figure 5A) before and after PA103 infection. Animals, which were either not subjected to serum transfer (data not shown) or treated with control sera, developed severe pneumonia and died within few days. In contrast, mice, which received serum from immune-(LD100-mAb166) animals survived better to PA103 infection as compared to those who received serum from non-protected-(LD0-mAb166) animals (Figures 5B and 5C). This result suggests that the humoral anti-PA103 response developed in animals resistant to secondary infection contributes to the protective anti-bacterial response.

**Figure 5:**
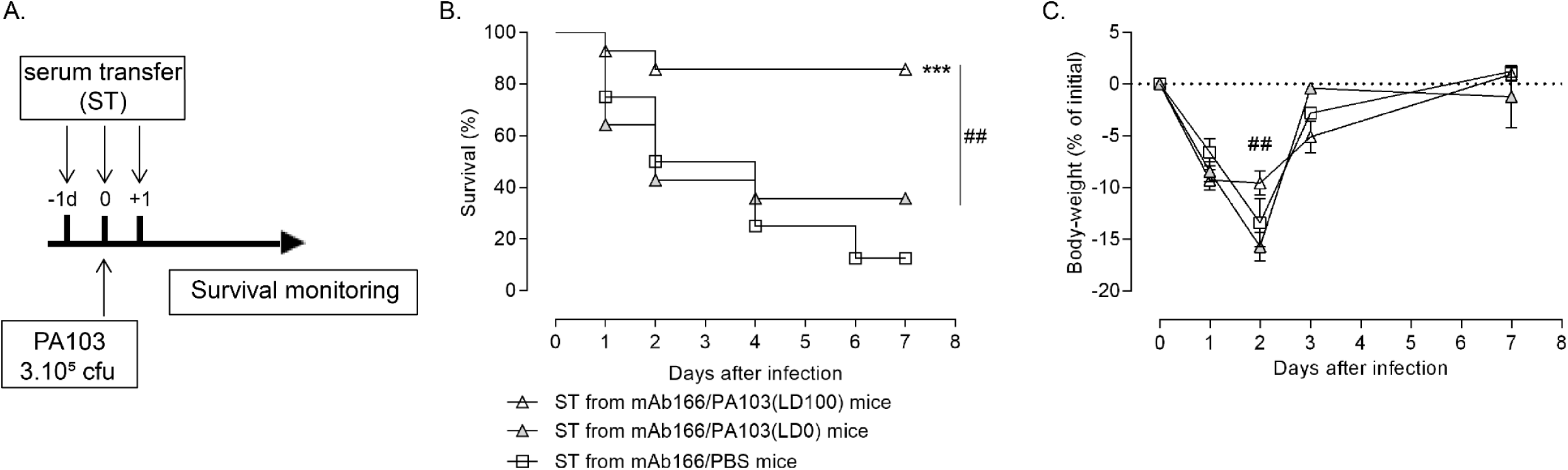
Serum from challenged immune mice protects against *P. aeruginosa* infection. (A) Serum from B6 mice that have been previously infected with *P. aeruginosa* (3.10^5^ cfu = LD100) and treated with mAb166 (100µg) were collected at D+28 after primary infection. Sera were injected intraperitoneally in naïve B6 mice at D-1, D0 and D+1 after an infection with *P. aeruginosa* (3.10^5^ cfu). Survival (B) and body-weight (C) was monitored over 7 days. The results correspond to 3 pooled, independent experiments (n=15 mice per group), ***: *p<*0.01 with a log-rank test comparing with mAb166/PBS group; ##: *p<*0.01 with a log-rank test comparing with mAb166/PA103(5.10^4^) group.

### Long-lasting humoral response induced by anti-bacterial Ab mediates cross-protection against *P. aeruginosa*

Although numerous *P. aeruginosa* antigens have been investigated as immunotherapy or vaccine candidates, their development have been hampered by the inability to achieve broad protection across different serotypes ^33^. Here, we investigated whether Ab-mediated long-term response in mice infected with PA103 (serotype O11) and treated with mAb166 may protect animals from a secondary infection by heterologous strains of *P. aeruginosa* (Figure 6A). We used CLJ1 and PA14 strains, which belong to the serogroups O12 and O10, respectively (Supplementary Table 1). PA14 express *pcrV*, while not CLJ1, but none of them are sensitive to mAb166 immunotherapy (Supplementary figure 5). While the long-term effect associated to the infection with PA103-3.10^5^cfu and the treatment with mAb166-100µg was not sufficient to protect animals against a lethal challenge with CLJ1 (Supplementary figure 7), it significantly protects individuals from a sublethal infection by CLJ1 or lethal infection by PA14 (Figure 6B). Interestingly, this protection was associated with the production of anti-CLJ1 or anti-PA14 IgG, which, even lower as compared to the production of anti-PA103 IgG, was significantly superior to controls (Figure 6C). These results suggest that prior lung infection rescued by passive inhaled immunotherapy induces a humoral protective immunity against *P. aeruginosa* pneumonia induced by heterologous strains.

**Figure 6:**
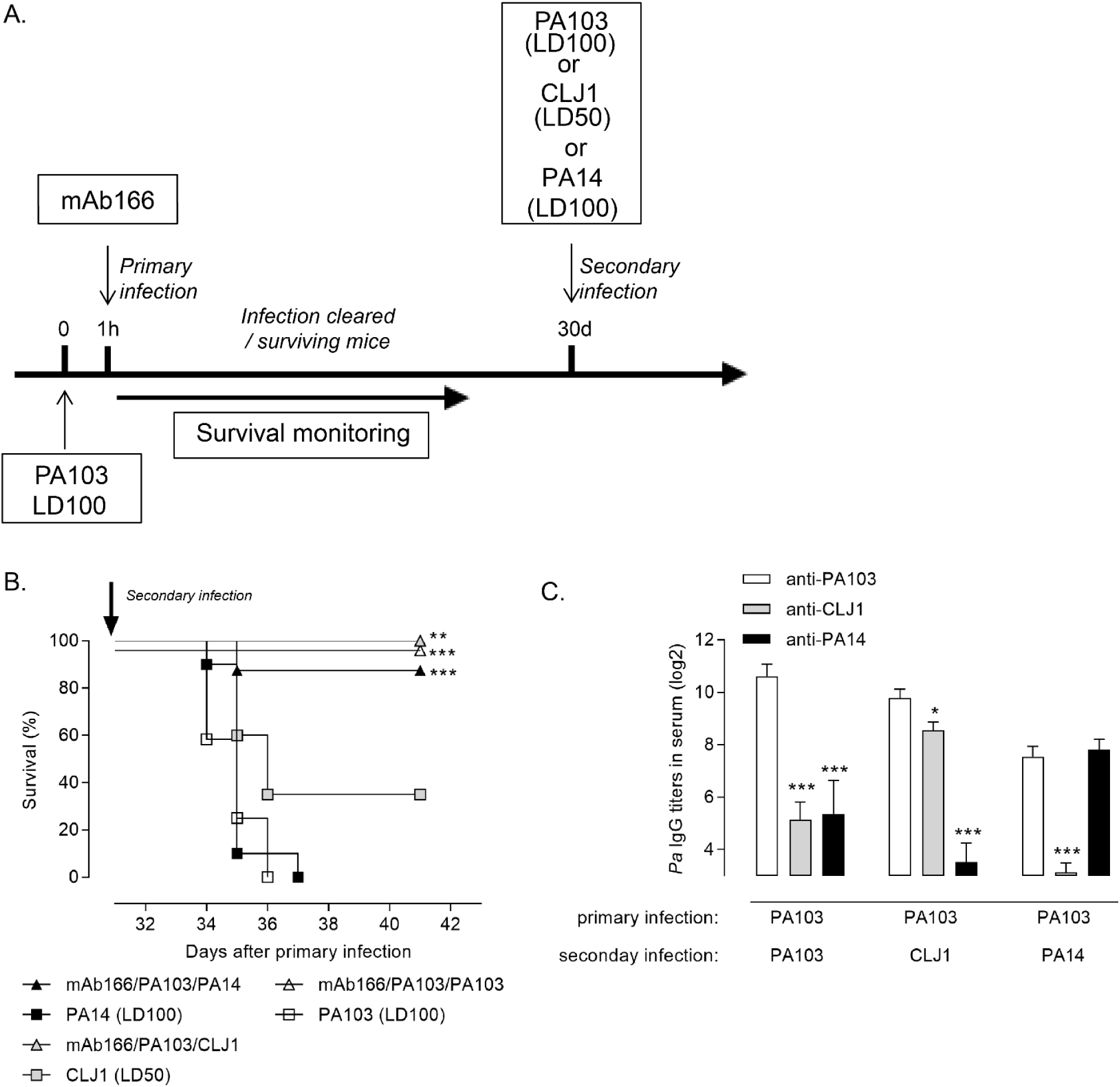
Long-lasting humoral response induced by anti-bacterial Ab mediates cross-protection against *P. aeruginosa*. (A) B6 mice were treated/infected as described in Figure 1C. At the challenge, mice were infected with either homologous strain PA103 WT (LD100), or heterologous strains CLJ1 (LD50) or PA14 (LD100). (B) Survival was monitored over the entire period. The results correspond to 3 pooled, independent experiments (n=10-36 mice per group), **: *p<*0.01; ***: *p<*0.001 with a log-rank test comparing with respective untreated groups. (C) Total anti-PA103 (white bars), anti-CLJ1 (black bars) or anti-PA14 (gray bars) IgG in serum was determined by ELISA at D+7 after secondary infection. The data are quoted as the mean values ± SEM. The results correspond to 3 pooled, independent experiments (n=5-10 mice per group).

## Discussion

Despite considerable advances in antimicrobial chemotherapy, treatment of *P. aeruginosa* infections has become challenging because of the increasing prevalence of intrinsic and acquired multi-drug resistance to antibiotics ^34-37^. Notably, *P. aeruginosa* has become the most common multi-drug resistant gram-negative bacteria causing pneumonia in hospitalized patients, associated with premature mortality ^38^. In addition, managing antibiotic-resistance infections to *P. aeruginosa* is costly, adding pressure on overburdened healthcare systems ^39^. Besides acute pneumonia, *P. aeruginosa* can also cause persistent infections in patients with chronic pulmonary diseases including cystic fibrosis (CF) ^40^, chronic obstructive pulmonary disease (COPD) ^41^ or non-CF bronchiectasis ^42^ and is also associated with recurrent infections in ICU patients ^43^. Anti-infectious Ab therapies have gained an important place in the therapeutic arsenal against infectious disease. As illustrated in the treatment of *Clostridium difficile* or HIV infections, they offer novel perspectives of addressing antimicrobial resistance and chronic/recurrent *P. aeruginosa* infections. Our group and others have demonstrated that the airways constitute an attractive and feasible alternative route for Abs delivery into the lungs ^19, 44-46^ enhancing their local concentration, limiting their passage into the systemic circulation and leading to a better anti-microbial protective response as compared to other delivery routes ^20, 22, 23, 47^. After mucosal delivery, antibody-mediated anti-infective response relies mostly on Fab-dependent neutralization of pathogen and recruitment of immune mediators via FcγR.

Using a robust and straightforward murine model of pneumonia mimicking the initial phases of lung infection during *P. aeruginosa* colonization of the airways, we demonstrated for the first time a long-lasting protection associated with mucosally-delivered Ab, protecting the individuals from a secondary infection. Recent clinical and preclinical studies have identified similar functions for Ab, which act as immunomodulators that could bridge innate, acquired, cellular and humoral immune responses. However, this Ab-mediated long-term effect was for instance limited to cancer or viral infection and after a systemic administration of the therapeutic agent ^3, 16^. Our results showed that this effect was not related to unspecific vaccination process (due to the bacteria), as the long-term immunity was dependent on the dose of the Ab and the size of bacteria inoculum while an antibiotic treatment was unable to promote protection against a secondary infection. Similarly, we observed that this effect was restricted to the presence of the antigen as demonstrated by the absence of protection after a secondary infection when the mice have been primary infected with a *P. aeruginosa* strain devoid of antigen expression. This indicates a critical role of immune-complexes (IC) between therapeutic Ab and bacteria expressing the cognate antigen, as suggested by the literature ^14, 48, 49, 50^, in the induction of a long-term protection. The duration and strength of this effect may be supported by the ability of antigen-presenting cells to internalize IC and promote presentation of epitopes from Ig-complexed antigens to T cells ^51, 52^.

In order to translate the observations of Ab-mediated long-term protection into human benefits, it appears essential to identify the molecular and cellular mechanisms supporting these long-term protective responses. During viral infections, Ab-mediated long-term protection relies on the induction of endogenous humoral response ^9, 10, 15, 53^, but not in cancer models ^14^. Here, we provided evidences that anti-*P. aeruginosa* humoral response is a critical component of the Ab-mediated long-term protection. First, anti-bacterial IgG and especially IgG3 are generated in a greater extent after primary infection in individuals that will subsequently survive to the challenge, with a direct effect on the control of the bacterial load. This subclass stimulated by a pro-Th1 inflammatory milieu ^54^ was demonstrated to actively contribute to the protection against other respiratory bacteria ^55^. IgG3 may also favor the emergence of a memory response after the challenge. Second, transferred anti-bacterial IgG enriched decomplemented serum provide protection against *P. aeruginosa* infection. Finally, both local and systemic B cells are increased in the context of long-term protection. Taken together, these results indicate that *P. aeruginosa*-specific memory humoral response induced by mucosal delivery of Ab mediates protection against repeated pulmonary infection. This is in accordance with previous evidences indicating an essential contribution of the humoral response to antiviral Ab long-term response ^49, 56^. Interestingly, in the context of cancer, the long-term antitumor protection induced by Ab therapy was independent from endogenous antibody response ^14^, suggesting distinct long-term protective immune responses depending on the inflammatory condition.

The precise mechanisms underlying the ability of memory humoral response to supply local and systemic IgG responses remains an important question to be addressed. *P. aeruginosa* infection leads to formation of tertiary lymphoid organs in the lungs, called inducible broncho-associated lymphoid tissues (iBALT) ^57^. These structures circumscribed the complex orchestration of B cells, innate immune cells and T cells ^58^ which are critical for the induction of recall responses against re-infecting pathogen ^59, 60^. Even if we provide some evidences of the involvement of specific subtypes of B cells in the induction of Ab-mediated the long-term protection, additional experiments are necessary to provide a complete picture of the cellular and molecular partners accounting for the induction of long-term anti-*P. aeruginosa* humoral response ^30, 61^.

*P. aeruginosa* is a highly versatile pathogen intrinsically associated with COPD or CF patients, acting as a colonizer that contributes to deterioration of lung function and fatal outcome ^62, 63^, or being also a causative agent of nosocomial pneumonia with a significantly higher mortality as compared to other pathogens ^64^ and even in re-admitted patients ^43^. In this context, understanding immune mechanisms of long-term protection against *P. aeruginosa* is critical to develop effective Ab-based therapies that are broadly protective against bacterial pneumonia. For instance, whether Ab-mediated long-lasting effects are solely directed at the Ab-targeted antigen or provide epitope spreading remains unclear and may depends on the pathological context ^3^. In this study, we observed that mice primarily immunized against PA103 strain were protected against secondary infections mediated by heterologous *P. aeruginosa* strains. However the broad-spectrum protection against *P. aeruginosa* induced by mucosal Ab is not as effective as specific immunity ^65^ to alter the course of pneumonia induced by heterologous strains and provide protection only against sublethal infection. The partial cross-protection may be due to a weaker endogenous humoral response generated against strains from different serotypes. This may be associated with lower IC immunoregulation, which is critical to improve cross-presentation and epitope spreading ^66, 67^.

In conclusion, mucosal delivery of Ab offers the best protection against *P. aeruginosa*, a high-priority classified pathogen for which there is currently no vaccine or Ab approved. It will allow rapid pathogen neutralization limiting its replication while providing an optimal host immune response to eliminate the pathogen and induce a long-lasting protection against secondary infections. This novel modality associated with anti-bacterial Ab may be of importance for the treatment of pathogen causing recurrent acute infections as well as chronic respiratory infections, especially in the context of antibiotic resistance.

## Materials and methods

### Mice and antibody

Adult male C57BL/6jrj (B6) mice (6-8 weeks old) were obtained from Janvier (Le Genest Saint-Isle, France). All mice were housed under specific-pathogen-free conditions at the PST “Animaleries” animal facility (Université de Tours, France) and had access to food and water *ad libitum*. All experiments complied with the French government’s ethical and animal experiment regulations (APAFIS#7608-2016111515206630).

mAb166 was generated from PTA-9180 hybridoma (LGC Standards, France) and supplied as sterile, non-pyrogenic, PBS solution under good manufacturing practice (BioXcell, USA).

### *P. aeruginosa* primary infection

Severa*l P. aeruginosa* strains were used in the study: PA103 (ATCC 33348) and PA103Δ*pcrV*, kindly provided by Pr. Teiji Sawa (Kyoto University, Japan), CLJ1 kindly provided by Dr. Ina Attree (CEA Grenoble, France) and PA14, kindly provided by Dr. Eric Morello (Tours University, France). The uniformity of the colonies was checked by plating on *Pseudomonas* isolation agar (PIA) plates. PA103 has been transformed by quadriparental mating by a mini-Tn7T transposon encoding allowing a constitutive expression of the *LuxCDABE* operon. Bacteria were prepared as previously described ^22^. Mice were anesthetized with isofluorane 4% and an operating otoscope fit with intubation specula was introduced to both maintain tongue retraction and visualize the glottis. A fiber optic wire threaded through a 20G catheter and connected to torch stylet (Harvard Apparatus, France) was inserted into the mouse trachea. Correct intubation was confirmed using lung inflation bulb test and 40µL of the bacterial solution was applied using an ultrafine pipette tip. Inoculum size for infections were confirmed by counting colony-forming unit (cfu) on PIA plates and were as follow: 5.10^4^ cfu (LD0 dose), or 5.10^5^ cfu (LD100, experiments with prophylactic intervention), or 3.10^5^ cfu (LD100, experiments with therapeutic setting) for PA103; 5.10^5^ cfu (non-lethal dose) for PA103Δ*pcrV*. Mortality and body weight of animals were monitored daily. In all experiments, moribund animals or animals with a weight loss of more than 20% were sacrificed for ethical reasons and considered as dead animals due to the infection.

### Antibody, antibiotic administration

mAb166 was either administered 2 hours before (prophylactic intervention) or 1 hour after the infection (therapeutic intervention), at the dose of 100µg or 50µg (suboptimal quantity) in 50µL of PBS using a Microsprayer® aerosolizer (Penn-Century, US) introduced orotracheally, as described in the previous section. Amikacin sulfate (Sigma, France), 25mg/kg was administered through the airways using the same protocol.

### *P. aeruginosa* secondary infection

Mice recovering from the primary infection were re-infected orotracheally as described above with PA103 – 5.10^5^ cfu (experiments with prophylactic intervention) or 3.10^5^ cfu (experiments with therapeutic intervention); CLJ1 – 1.10^6^ cfu (LD50) or PA14 – 1.10^6^ cfu (LD100), 30 days after the primary infection.

### Blood sampling and adoptive serum transfer experiments

Blood samples were collected every 3 days after the primary infection and at day 1, 5 and 7 after the secondary infection to assess endogenous anti-*P. aeruginosa* IgG concentrations. At day 28 after the primary infection, mice were sacrificed and sera were collected, pooled, heat-inactivated (56°C) for 30 minutes and diluted 1:6 in PBS. For serum transfer experiments, age-matched mice were administered intraperitoneally 100µL of 1/6 fold diluted serum in 1X PBS, on days -1, 0, and +1 post-infection with a LD100 of PA103 (3.10e5 cfu).

### Broncho-alveolar lavage, organ sampling, bacterial load assay

Broncho-alveolar lavage fluid (BALF) was collected after *P. aeruginosa* infection by introducing a catheter into the trachea under deep pentobarbital anesthesia and washing sequentially the lung with 1 × 0.5 mL and 2 × 1mL of 1X PBS at room temperature. The lavage fluid was centrifuged at 400 g for 10 min at 4°C and the supernatant of the first lavage was stored at -20°C for analysis. The cell pellet was resuspended in PBS, counted in a haemocytometer chamber and used for subsequent analysis. Spleen and lungs were perfused with 10mL of PBS1X and harvested in GentleMACS C tubes (Miltenyi Biotec, Germany) containing 2mL of RPMI medium (Invitrogen, France) for flow cytometry or GentleMACS M tubes (Miltenyi Biotec, Germany) containing 2mL of 1X PBS for microbiology assay.

Bacterial load in lung homogenates or BAL (before centrifugation) was determined by plating tenfold serial dilutions on PIA-agar. Plates were incubated at 37°C in a 5% CO_2_ atmosphere, and the cfu were counted after 24 hours.

### Preparation of pulmonary and spleen immune cells

Lung and spleen homogenates were prepared using a GentleMACS tissue homogenizer (Miltenyi Biotec, Germany). Lung pieces were then digested in a medium containing 125µg/mL of Liberase (Roche, France) and 100µg/mL of DNAse I (Sigma, France) for 30min at 37°C under gentle agitation. After washes, contaminating erythrocytes were lysed using Hybri-Max® lysis buffer (Sigma, France) according to manufacturer’s instructions. Samples were sequentially filtered over 100µm and 40µm nylon mesh. After final wash, cell pellets were resuspended in PBS containing 2 % FCS, 2mM EDTA and 1X murine Fc-block (Becton Dickinson, France) – described elsewhere as FACS buffer.

### Flow cytometry

Cells were incubated in FACS buffer for 20min at 4°C. Then, cells were stained in FACS buffer for 20 min at 4°C with appropriate dilutions of following antibodies: CD45-APC-Cy7 (30-F11), CD3 PerCP-Cy5.5 (17A2), CD19 V450 (1D3), IgM PE (RMM-1), CD21 APC (7E9), CD11b PercP (M1/70), and Ly6G FITC (1A8) from Biolegend and Siglec-F BV421 (E50-2440) from BD. Dead cells were stained with the LIVE/DEAD Fixable Aqua Dead Cell Staining kit (ThermofisherScientific) and acquired on a MACS Quant (Miltenyi Biotec) cytometer. Analyses were performed using Venturi-One software (Applied Cytometry; UK)

### Cytokines, mAb166 and IgG measurements

Pa-specific IgG, IgG1 and IgG3 titers, in serum and BALF levels were measured by sandwich ELISA. Briefly, high binding Immulon 96-well plates (ThermoFischer Scientific, France) were coated with 0.5µg/mL of PA103, PA14 or CLJ1 lysates, prepared from overnight cultures that were then sonicated, and diluted in bicarbonate buffer. The plates were then washed and blocked with 1%BSA-PBS. Serum or BALF samples were incubated for 2 h. A biotin-conjugated goat anti-rat IgG antibody (Biolegend, France) was added for 2 h. After a washing step, peroxidase-conjugated streptavidin (R&D) was added for 20min. Between each step, plates were thoroughly washed in 0.05%Tween20-PBS. Tetramethylbenzidine was used as a substrate, and the absorbance was measured at 450 nm using a microplate reader (Tecan, Switzerland). The titer was calculated by binary logarithm regression as the reciprocal dilution of the sample, where the extinction was 2-foldthe background extinction. Similar procedure was used to quantify mAb166 in serum and BALF using recombinant pcrV (0.5µg/mL) as coating antigen. Before assay, antibody-antigen dissociation was performed, using acetic acid, as previously described ^22^. Cytokines concentrations in BALF were assessed using a Bio-plex magnetic bead assay (Luminex, Biotechne, France) and analyzed using a Bio-plex 200 workstation (Biorad, France).

### Statistical analysis

Statistical evaluation of differences between the experimental groups was determined by using one-way analysis of variance (ANOVA) followed by a Newman-Keuls post-test (which allows comparison of all pairs of groups). Log-rank test was used for survival analysis. Student’s t-test was used for comparison between two groups and paired t-test was used when comparing the same individual before and after challenge. All tests were performed with GraphPad Prism, Version 6 for Windows (GraphPad Software Inc., San Diego, CA, USA; www.graphpad.com). All data are presented as mean +/- standard error of the mean (SEM). A p value <0.05 was considered significant.

## Acknowledgements

This work was supported by a public grant overseen by the French National Research Agency (ANR) as part of the “Investissements d’Avenir” program (LabEx MAbImprove, ANR-10-LABX-53-01). AP is funded by a fellowship from ANR-10-LABX-53-01. Additional fundings were provided by Region Centre-Val-de-Loire (Novantinh Program) and C-VALO (Infinhitim Program).

## Author Contributions

TS and NHV conceived the study. All authors substantially contributed to the acquisition, analysis or interpretation of data. CP provided expertise in the analysis of the immune response. AP, TS and NHV contributed to manuscript drafting, revising and critically reviewing. All authors approved the final version of this manuscript to be published.

## Figure Legends

<u>Supplementary Figure 1: Kinetics of free and total mAb166 in the serum and the airways after PA103 infection.</u>

B6 mice were treated/infected as described in Figure 1C. The free (A, B) and total (C, D) mAb166 concentrations in serum (A, C) and in BALF (B, D) were determined by ELISA at days 1, 3, 7, 10 and 14 after primary infection. The data are quoted as the mean values ± SEM. The results correspond to 4 pooled, independent experiments (n=5 mice per time-points).

<u>Supplementary Figure 2: Morbidity and mortality of isogenic</u> *<u>pcrV</u>* <u>mutant PA103Δ</u>*<u>pcrV</u>* <u>strain.</u>

(A) B6 mice were infected as described in Figure 1A. (B) Body-weight loss and (C) survival were monitored over 7 days. The data are quoted as the mean values ± SEM. The results correspond to 2 pooled, independent experiments (n=10 mice per group). **: *p<*0.01 with a log-rank test. **: *p<*0.01, with a two-way ANOVA followed by a Bonferonni’s post-test.

<u>Supplementary Figure 3: Morbidity and mortality of PA103 according to the inoculum size.</u>

(A) B6 mice were infected as described in Figure 1A. (B) Body-weight loss and (C) survival were monitored over 7 days. The data are quoted as the mean values ± SEM. The results correspond to 3 pooled, independent experiments (n=8-9 mice per group). ***: *p<*0.001 with a log-rank test. **: *p<*0.01; ***: *p<*0.001 with a two-way ANOVA followed by a Bonferonni’s post-test.

<u>Supplementary Figure 4: Prophylactic inhaled administration of anti-bacterial Ab is associated with a sustained anti-</u>*<u>P. aeruginosa</u>* <u>humoral response.</u>

B6 mice were treated/infected as described in Figure 1A. (A) The concentration of total anti-PA103 IgG in serum was determined by ELISA at days 7, 14, 21 and 30 after the primary infection. The data are quoted as the mean values ± SEM. The results correspond to 9 pooled, independent experiments (n=5-56 mice per time-points), **: *p<*0.01; ***: *p<*0.001 with two-way ANOVA followed by a Bonferonni’s post-test. (B) The concentration of total anti-PA103 IgG in serum was determined by ELISA at days 1, 5 and 7 after the secondary infection. The data are quoted as the mean values ± SEM. The results correspond to 3 pooled, independent experiments (n=15 mice per time-points). (C) The concentration of total anti-PA103 IgG in serum was determined by ELISA at days -1, +5 after the secondary infection. The data are quoted as individual values. The results correspond to 5 pooled, independent experiments (n=21 mice per time-points), ***: *p<*0.001; with a paired t-test. Frequency (left axis) and total number (right axis) of B cells (CD45+ CD3-CD19+ cells) was determined in the lungs (E) and the spleen (F) of animals at D+1 after secondary infection. The data are quoted as the mean values ± SEM. The results correspond to 2 pooled, independent experiments (n=8 mice per group), *: *p<*0.05 with a parametric t-test.

<u>Supplementary Figure 5: Primary infection with heterologous</u> *<u>P. aeruginosa</u>* <u>strains is not rescued by mAb166.</u>

(A) B6 mice were infected as described in Figure 1C. (B) Survival was monitored over 7 days. The data are quoted as the mean values ± SEM. The results correspond to 3 pooled, independent experiments (n=10-38 mice per group).

<u>Supplementary Figure 6: Long-term protection induced by inhaled mAb166 is insufficient against LD100 of</u> *<u>P. aeruginosa</u>* <u>CLJ1 strains.</u>

(A) B6 mice were infected as described in Figure 1C. (B) Survival was monitored over 7 days. The data are quoted as the mean values ± SEM. The results correspond to a pool of three independent experiments (n=10-15 mice per group).

