## Supplementary figures for "Mucosal administration of anti-bacterial antibody provides long-term cross-protection against *Pseudomonas aeruginosa* respiratory infection"

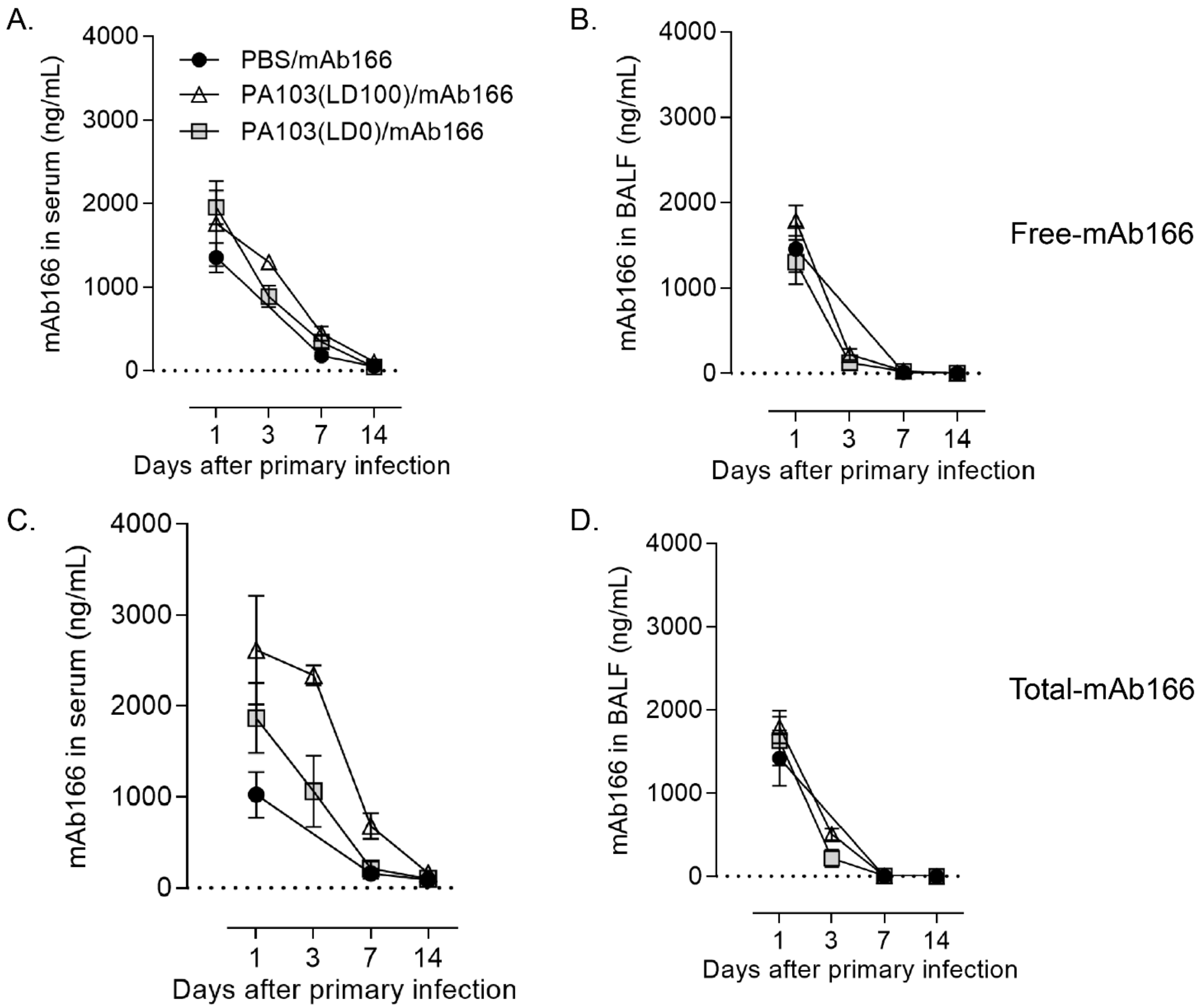


Supplementary Figure 1:

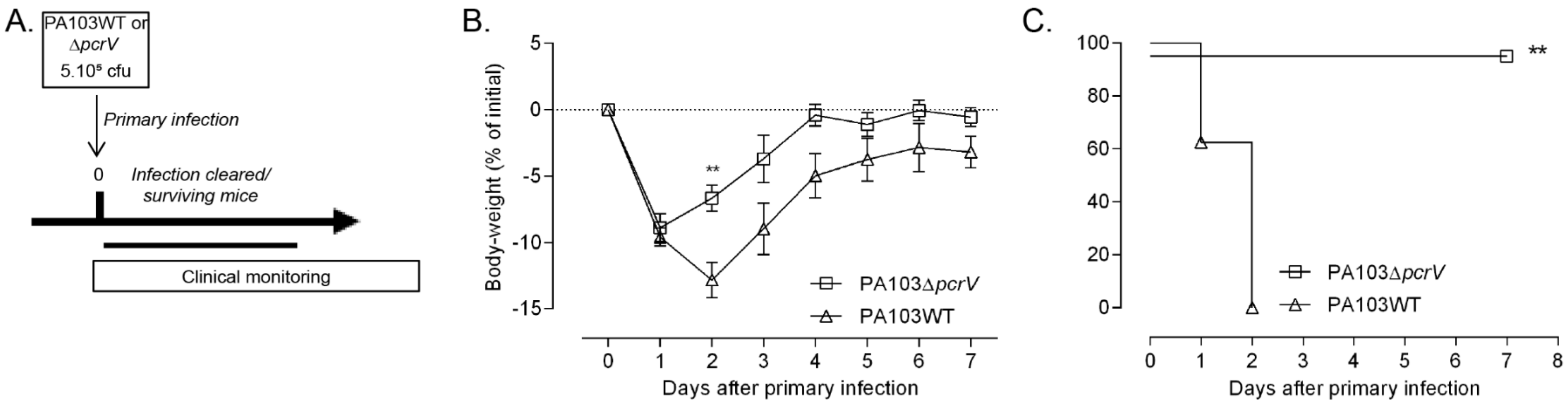


Supplementary Figure 2:

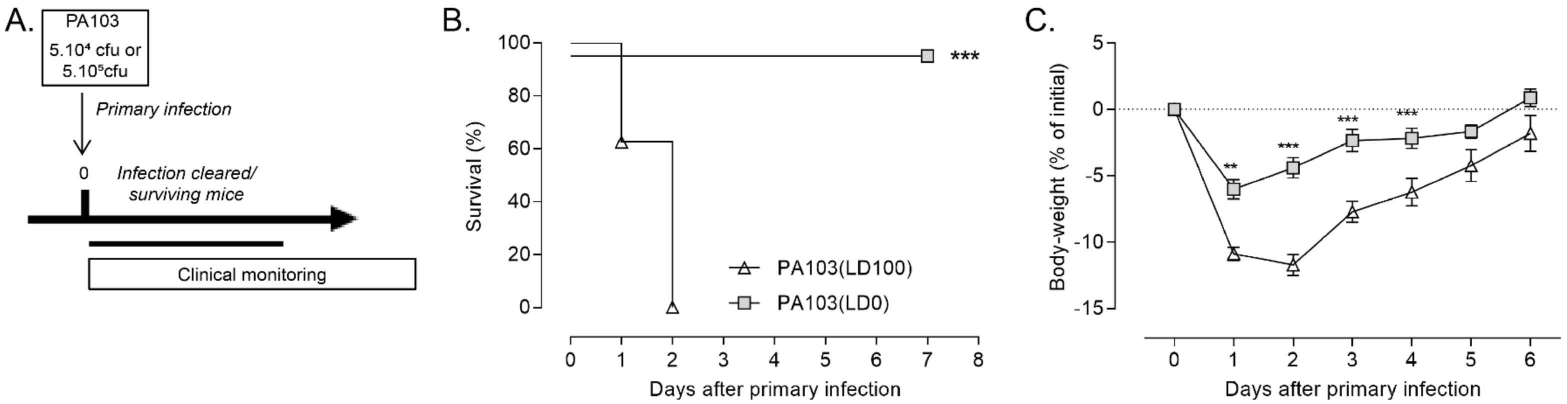


Supplementary Figure 3:

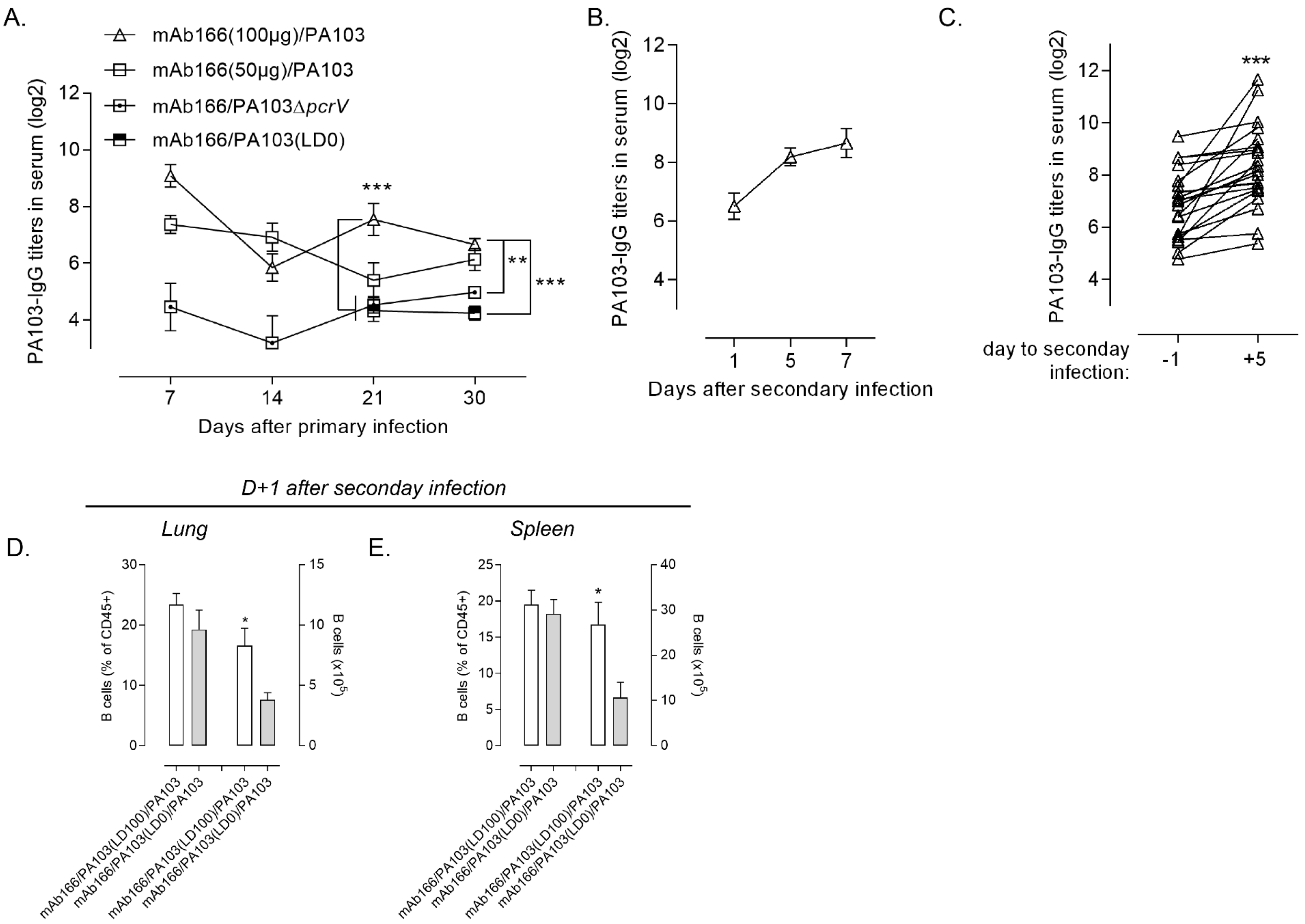


Supplementary Figure 4:

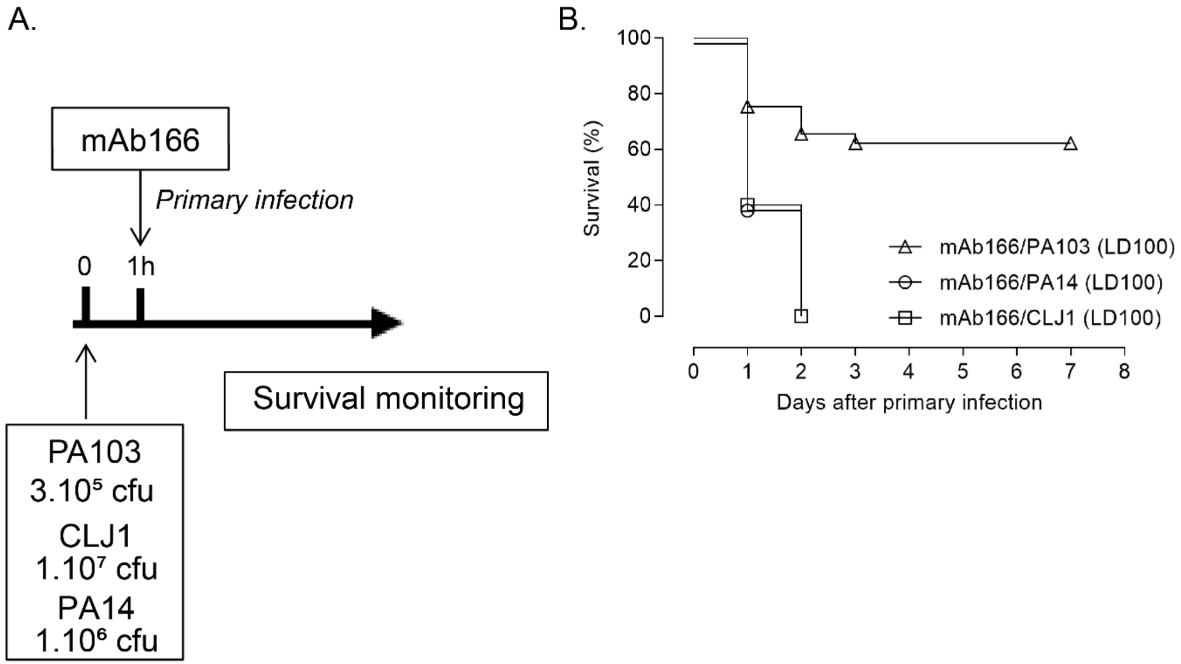


Supplementary Figure 5:

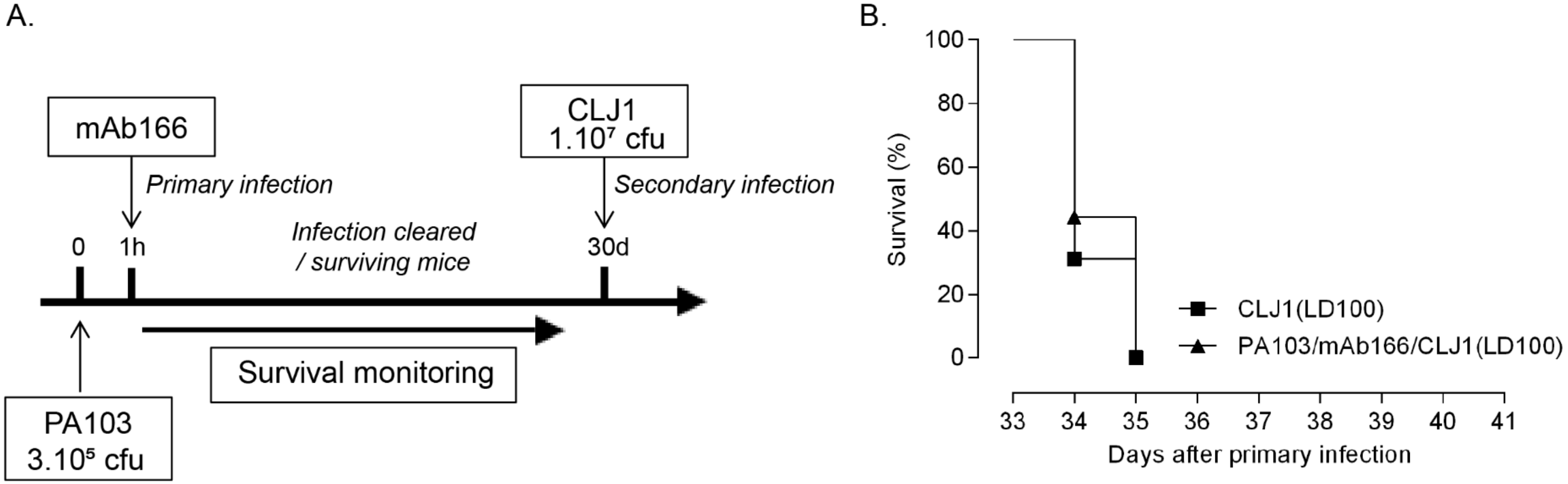


Supplementary Figure 6:

Supplementary table 1: Characteristics of *P. aeruginosa* strains used in this study

| Name | Serotype | *pcrV* |
|---|---|---|
| *PA103* | O11 | + |
| *CLJ1* | O12 | - |
| *PA14* | O10 | + |

Supplementary Table 1:
